# Retracing Schwann cell developmental transitions in embryonic dissociated DRG/Schwann cell cocultures in mice

**DOI:** 10.1101/2020.07.31.231258

**Authors:** Venkat Krishnan Sundaram, Tatiana El Jalkh, Rasha Barakat, Camille Julie Isabelle Fernandez, Charbel Massaad, Julien Grenier

**Affiliations:** Université de Paris, INSERM UMRS 1124, Faculty of Basic and Biomedical Sciences, Paris, France; Université de Paris, INSERM UMRS 1016, Institut Cochin, Paris, France; Lebanese University, EC2M, Faculty of Sciences II, Fanar, Lebanon

**Keywords:** Schwann cell development, Dissociated DRG/SC cocultures, Schwann Cell Precursors, immature Schwann Cells, myelinating Schwann cells

## Abstract

Embryonic Dissociated Dorsal Root Ganglia cultures are often used to investigate the role of novel molecular pathways or drugs in Schwann cell development and myelination. These cultures largely recapitulate the order of cellular and molecular events that occur in Schwann cells of embryonic nerves. However, the timing of Schwann cell developmental transitions, notably the transition from Schwann Cell Precursors to immature Schwann cells and then to myelinating Schwann cells, has not been estimated so far in this culture system. In this study, we determined the expression profiles of Schwann cell developmental genes during the first week of culture and then compared our data to the expression profiles of these genes in developing spinal nerves. This helped in identifying that Schwann Cell Precursors transition into immature Schwann Cells between the 5^th^ and 7^th^ day *in vitro*. Furthermore, we also investigated the transition of immature cells into pro-myelinating and myelinating Schwann cells upon the induction of myelination *in vitro*. Our results suggest that Schwann cell differentiation beyond the immature stage can be observed as early as 4 days post the induction of myelination in cocultures. Finally, we compared the myelinating potential of coculture-derived Schwann cell monocultures to cultures established from neonatal sciatic nerves and found that both these cultures system exhibit similar myelinating phenotypes. In effect, our results allow for a better understanding and interpretation of coculture experiments especially in studies that aim to elucidate the role of a novel actor in Schwann Cell development and myelination.

## Introduction

Dissociated Dorsal Root Ganglia (DRG) cultures from mouse embryos have long been utilized as a resourceful model for exploring the nuances of Schwann cell development *in vitro* (Taveggia and Bolino, 2018). The co-culture system provides a solid experimental framework to study different aspects of Schwann cell development such as proliferation, migration, differentiation and myelination of axons (Päiväläinen et al., 2008; Taveggia and Bolino, 2018). Furthermore, it recapitulates the different aspects of Schwann cell development that is observed *in vivo*. Hence, dissociated DRG cultures form an indispensable part of studies that aim to understand the role of a novel actor in Schwann cell development and differentiation.

It is well known that temporal differences exist between Schwann cell development in Dissociated DRG/SC coculture *in vitro*, and in developing spinal nerves *in vivo*. In developing spinal nerves of mice, Neural Crest Cells (NCC), destined to a glial fate differentiate into Schwann Cell Precursors (SCP) and appear in the DRGs at around E11 (Jacob, 2015). Then, the SCPs start migrating on nascent axons between E12.5 and E13.5 to populate their peripheral targets. However, at around E15.5 in mice, SCPs undergo a transition into immature Schwann cells (iSC) that further differentiate into either myelinating or non-myelinating Schwann cells, perinatally (Monk et al., 2015; Fledrich et al., 2019; Jessen and Mirsky, 2019).

Nevertheless, these observations cannot be used to extrapolate the timing of Schwann cell developmental transitions *in vitro* because of certain technical issues. Firstly, the DRGs are dissected from mouse embryos towards the end of the 2^nd^ week of gestation (E12.5 or E13.5). At this stage, SCP *in vivo* have already started departing from the DRGs and begun migrating on developing axons (Jessen et al., 1994; Jessen and Mirsky, 2005). However, once dissected and dissociated, the E13.5 DRG cells give rise to sensory neurons and Schwann cell precursors once again *in vitro* (Ratner et al., 2005; Kim and Maurel, 2009; Kim and Kim, 2018). This is rendered possible because of a reservoir of sensory neurons and SCP located inside the DRGs that repopulate the culture **(Figure 1)**. Therefore, a significant portion of *in vivo* developmental events are repeated in cell culture albeit with a phase difference. Our objective in this study is to better understand these temporal differences in cocultures in an effort to provide a better experimental and inferential framework.

**Figure 1:**
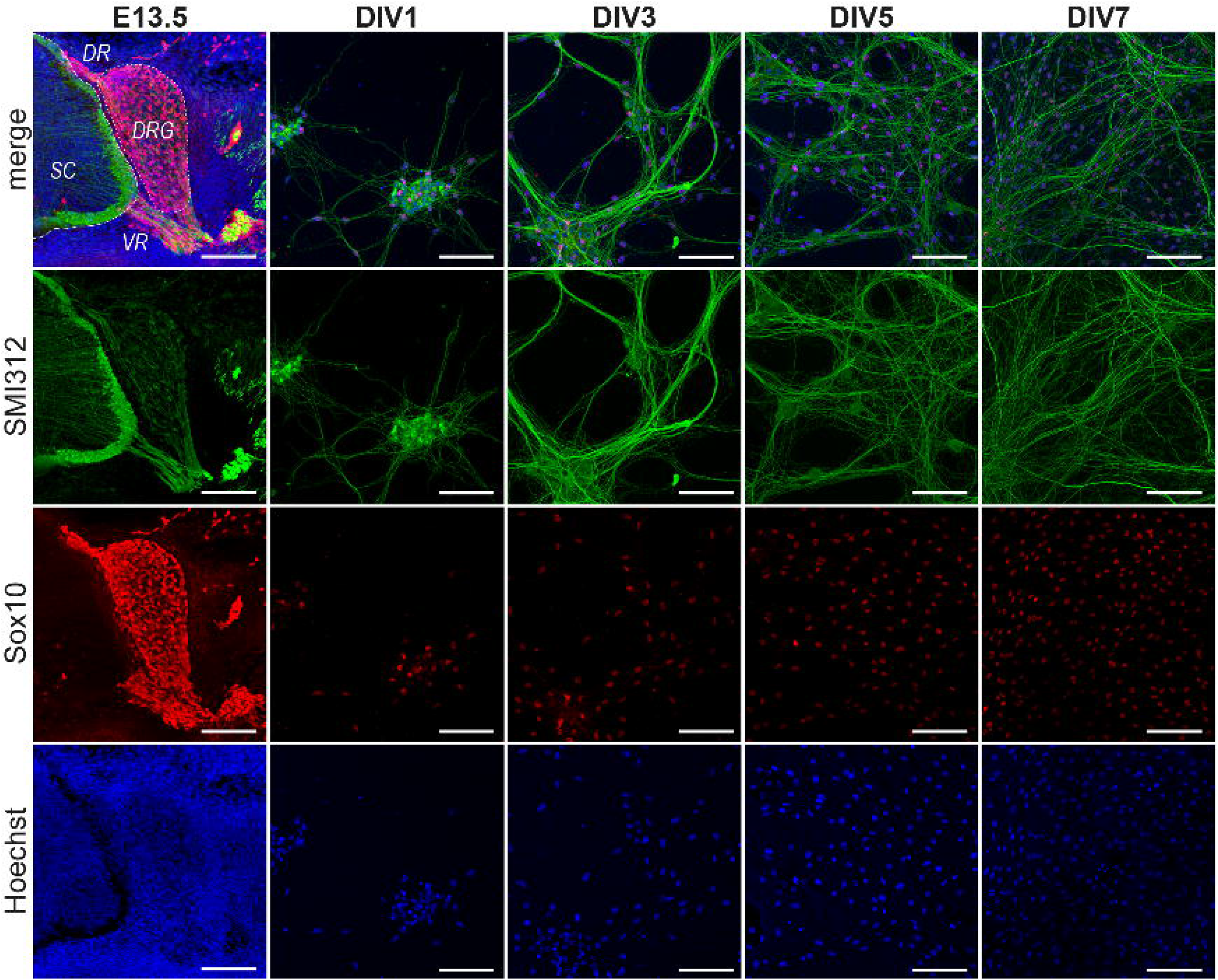
Progression of DRG/SC cocultures between Days in Vitro (DIV)1 & DIV7. DRGs were dissociated and cultured from E13.5 embryos. The DRGs at the time of dissections contained SMI312+ neurons and Sox10+ glial cells. The cells sparsely populated the culture at DIV1. In the next 5 days, the neuron-SC network expanded and established a well-connected network by DIV 7 containing Sox10+ SC situated on top of SMI312+ axonal extensions. Images shown here are representative images chosen arbitrarily. Although the area covered by axons increases progressively between DIV1 and DIV7, not all regions of the culture are equally dense at DIV7. SMI312 – Neuronal Marker, Sox10 – SC lineage marker, Hoechst – Nuclear Staining. SC: Spinal Cord, DR: Dorsal root, VR: Ventral root, DRG: Dorsal Root Ganglia. Scale bar = 100μm

To this end, we first delineated the mRNA expression profiles of the genes expressed in SCP (*Dhh, Mpz, Cnp, Plp, Mbp, Cad19 and Tfap2α*) and iSC (*Krox20*). We then compared them to their profiles *in vivo* described in previous high-throughput studies conducted on embryonic peripheral nerves that provide extensive data on the differential expression of genes during Schwann Cell developmental transitions (Buchstaller et al., 2004; D’Antonio et al., 2006) This analysis helped us in determining the exact time window when SCP transition into iSC in Dissociated DRG/SC cocultures. Furthermore, we also verified that iSC at DIV7 transition into pro-myelinating (pro-mSC) and myelinating Schwann cells (mSC) upon the induction of myelination by the addition of Ascorbic Acid (AA) to DIV7 cocultures. Finally, we investigated if SC monocultures established from DIV7 cocultures possess the same myelinating potential as monocultures established from neonatal sciatic nerves.

Taken together, our results show that SCP transition into iSC between DIV5 and DIV7 in cocultures. Furthermore, we have also observed that the iSC/mSC transition in co-cultures occurs as early as 4 days post Ascorbic acid treatment. As for Schwann cell monocultures, SC obtained from DIV7 cocultures differentiate into mSC similar to SC isolated from neonatal sciatic nerves, further suggesting that the cells at DIV7 are comparable to neonatal Schwann cells in culture. In conclusion, our data provides a holistic understanding of the Schwann Cell Precursor/immature Schwann Cell/myelinating Schwann Cell transition in embryonic DRG/SC cocultures which is crucial for designing rigorous *in vitro* assays to study Schwann cell embryonic development and embryonic phenotypes of different Schwann cell mutants.

## Materials and methods

### Animals and Tissue Harvesting

Timed pregnant C57Bl6/J mice at E13.5 were purchased from Janvier Labs. The pregnant mice were first anesthetized with isoflurane and sacrificed using cervical dislocation. Embryos were surgically removed and placed in ice-cold L-15 media. DRGs were harvested from these embryos based on existing protocols (Kim and Kim, 2018; Taveggia and Bolino, 2018). All aspects of animal care and animal experimentation were performed in accordance with the relevant guidelines and regulations of INSERM and Université de Paris (authorization APAFIS#7405-2016092216181520).

### Dissociated DRG/SC cocultures

A total of 40 DRGs were harvested from each embryo. DRGs were then trypsinized (0.25% Trypsin in HBSS1X) for 30min at 37°C. Trypsinization was stopped using L-15 media containing 10% Horse Serum (Gibco). DRGs were then spun down at 1500 rpm for 5min. The supernatant was removed, and the tissues were resuspended in *DRG plating medium* (refer to *Media Compositions* in Supplementary Methods). The tissues were then triturated 10-20 times using flamed Pasteur pipettes until a homogenous cell suspension was obtained. For each time point (DIV1, DIV3, DIV5, DIV7), dissociated DRGs were plated on 12 well plates containing 14mm coverslips coated with Poly L Lysine (Sigma) and Collagen (R&D Systems). 40 DRGs from each embryo were plated into 8 wells at approximately 5 dissociated DRGs per well. 2 wells were assigned to each time point. The cells were first plated with DRG plating medium for 16h. The following day the medium was replaced with Supplemented Neurobasal medium to promote neurite growth and Schwann cell proliferation for a period of 7 days. Media was changed every 48 hours. Myelination was induced at DIV7 by changing the media to DRG plating medium supplemented with 50μg/mL Ascorbic Acid (Sigma).

### Schwann cell monocultures

SC monocultures were established using previously detailed protocols (Kim and Kim, 2018). Briefly, dissociated DRGs were obtained from E13.5 and were cultured on uncoated 35mm petri dishes (approx. 40 DRGs/Embryo) as explained above. At DIV7, the neurite network (neurons + Schwann cells) was mechanically lifted from the plate using a sterile 27 ^L1/2^ G needle. The network was then enzymatically digested (0.25% Trypsin, 0.1% Collagenase in HBSS1X) for 30 min at 37°C. Digestion was stopped by the addition of *Schwann Cell Plating Media* and the cell suspension was centrifuged at 1500rpm for 5min. The pellet was then triturated 5 – 6 times using a 1mL pipette tip. To obtain highly pure cultures without contaminating fibroblasts, the cell suspension was subjected to immunopanning to remove Thy1.2+ve fibroblasts as described elsewhere (Lutz, 2014). About 500000 Schwann cells were obtained from each embryo after immunopanning and the cells were plated on 14mm Poly L Lysine coated glass coverslips at a density of 50000 cells/coverslip. Schwann cells were expanded using defined *Schwann Cell Proliferation Media* (see supplementary methods) for 48h. To induce differentiation, cell cultures were treated with *Schwann Cell Differentiation Media* (Proliferation media without Forskolin but supplemented with 1mM dbcAMP) for a period of 48h.

### Immunohistochemistry

E13.5 and E16.5 embryos were surgically removed from the pregnant mouse and placed on ice cold L-15 media. The head, the thoracic region along with the ventral internal organs and the tail were dissected. The lumbar region along with the hindlimbs were fixed overnight with 4% PFA at 4°C. The following day, the embryos were extensively washed with PBS1X and incubated overnight in Antigen Retrieval Buffer (10mM Sodium Citrate, 0.05% Tween20, pH 6.0) at 4°C. The following day, the samples were boiled in the antigen retrieval buffer for 5 min and immediately placed in ice cold 30% sucrose solution. The samples were then dehydrated in Sucrose overnight at 4°C. The following day, samples were embedded in 4% Agarose and placed on the vibratome such that the caudal aspect of the embryo was facing the chuck, the rostral aspect was facing upwards, and the lateral aspect was facing the blade. 50μm serial transverse sections of the lumbar region was made and transferred to a 12 well plate containing PBS1X. Sections were quickly washed in PBS1X and then stored at −20°C in a cryoprotectant (30% Glycerol, 30% Ethylene Glycol in PBS1X) until immunostaining.

Immunostaining was performed by washing the sections first with PBS1X followed by incubation in 0.1M Glycine for 1 hour. Sections were permeabilized and blocked with blocking buffer (0.5% Triton X100, 0.1% Tween20, 2% BSA and 5% Normal Donkey Serum) for 1 hour followed by incubation with primary antibodies against Neurofilament SMI312, Sox10 and Tfap2α for 36h at 4°C (refer to *Supplementary Methods* for primary and secondary antibody references and concentrations). Sections were then washed thrice (1h per wash) in PBS1X containing 0.1% Tween20 and were then incubated with the corresponding secondary antibodies for 1h at RT in the dark. The sections were then washed, and nuclei were stained using Hoechst 33342 dye. Samples were then mounted on slides using Permaflour (Thermo Fisher Scientific) and stored at 4°C till confocal imaging.

### Immunocytochemistry

DRG/SC cocultures and Schwann cell monocultures at different conditions were first fixed with 4% PFA at RT for 30 mins. The coverslips were then washed with PBS1X and stored at −20°C in a cryoprotectant until immunostaining.

Immunostaining was first performed by washing the coverslips first with PBS1X followed by incubation in Antigen Retrieval Buffer (refer to *Immunohistochemistry*) preheated to 95°C for 3min. Samples were then washed with PBS1X and incubated in 0.1M Glycine solution for 30 minutes followed by permeabilization (0.25% Triton X100 0.1% Tween 20 in PBS1X, 20 min at RT) and blocking (2% BSA, 0.1% Tween 20, 10% Normal Donkey Serum, 1h at RT). Coverslips were then incubated with primary antibodies against, Neurofilament SMI312, Sox10, Tfap2α, Oct6, Krox20 and Ki67 overnight at 4°C (refer to *Supplementary Methods* for primary and secondary antibody references and concentrations). The following day, the coverslips were washed thrice with PBS1X (10 min per wash) and incubated with corresponding secondary antibodies for 1h at RT in the dark. Samples were then washed, and nuclei were stained with Hoechst 33342 dye. Samples were then mounted on slides using Permafluor (Thermo Fisher Scientific) and stored at 4°C till confocal imaging.

### Imaging and Image analysis

Confocal imaging of tissues sections and coverslips were performed on the LSM710 microscope. Images were obtained as z-stacks and analyzed in ImageJ. For each experimental condition, 3 – 4 biological replicates (embryos) and 3 technical replicates/biological replicate were analyzed. Cell counting was performed on z-projections (Max intensity) using the Analyze particles function after thresholding the images. Data was exported to Microsoft excel and graphs were plotted using Prism v8.0

### Total RNA isolation

Total RNA was extracted from each sample using 1mL of TRIzol reagent (Ambion Life Technologies 15596018) on ice as described in the manufacturer’s instructions with slight modifications. Briefly, 100% Ethanol was substituted for Isopropanol to reduce the precipitation of salts. Also, RNA precipitation was carried out overnight at −20°C in the presence of glycogen. The following day, precipitated RNA was pelleted by centrifugation and washed at least 3 times with 70% Ethanol to eliminate any residual contamination. Tubes were then spin dried in vacuum for 5 minutes and RNA was resuspended in 20μL of RNA resuspension buffer containing 0.1mM EDTA, pH 8. RNA was then stored at −80°C till RTqPCR.

### RNA quality, integrity and assay

RNA quantity was assayed using UV spectrophotometry on Nanodrop One (Thermo Scientific). Optical density absorption ratios A260/A280 & A260/A230 of the samples were above 1.8 and 1.5, respectively. The yield (mean ± SD) for each time point is as follows: DIV1 (26.74 ± 2.57 ng/μL), DIV3 (61.3 ± 8.01 ng/μL), DIV5 (51.86 ± 10.8 ng/μL), and DIV7 (77.34 ± 24.04 ng/μL). The extraction protocol used in the study was also validated using Agilent Bioanalyzer (RIN value 9.0 and above).

### RTqPCR

250ng of Total RNA was reverse transcribed with Random Primers (Promega C1181) and MMLV Reverse Transcriptase (Sigma M1302) according to prescribed protocols. Quantitative Real time PCR (qPCR) was performed using Absolute SYBR ROX 2X qPCR mix (Thermo AB1162B) as a fluorescent detection dye. All reactions were carried out in a final volume of 7μl in 384 well plates with 300 nM gene specific primers, around 3.5ng of cDNA (at 100% RT efficiency) and 1X SYBR Master Mix in each well. Each reaction was performed in triplicates. All qPCR experiments were performed on BioRad CFX384 with a No-Template-Control (NTC) to check for primer dimers and a No-RT-Control (NRT) to check for any genomic DNA contamination.

### Primer design and efficiency

All primers used in the study were designed using the Primer 3 plus software (https://primer3plus.com/cgi-bin/dev/primer3plus.cgi). Splice variants and the protein coding sequence of the genes were identified using the Ensembl database (www.ensembl.org). Constitutively expressed exons among all splice variants were then identified using the ExonMine database (https://imm.medicina.ulisboa.pt/group/exonmine/ack.html) (Mollet et al., 2010). Primer sequences that generated amplicons spanning two constitutively expressed exons were then designed using the Primer 3 plus software. For detailed information on Primer sequences used in the study, refer to the Supplementary Methods. The amplification efficiencies of primers were calculated using serial dilution of cDNA molecules. Briefly, cDNA preparations from all the time points were pooled and serially diluted three times by a factor of 10. qPCR was then performed using these dilutions and the results were plotted as a standard curve against the respective concentrations of cDNA. Amplification efficiency (E) was calculated by linear regression of standard curves using the following equation:

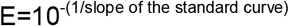

Primer pairs that exhibited a theoretical Amplification Efficiency (E) of 1.9 to 2.1 (95% - 105%) and an R^2^ value (Determination Coefficient) of 0.98 and above were chosen for this study.

### qPCR statistical analysis and Data Visualization

qPCR readouts were analyzed in Precision Melt Analysis Software v1.2. The amplicons were subjected to Melt Curve analysis and were verified for a single dissociation peak at a Melting Temperature (T_m_) > 75°C as expected from the primer constructs. The Cq data was exported to Microsoft Excel for further calculations. Each biological sample had 3 technical replicates thereby generating 3 individual Cq values. The arithmetic mean of the triplicates was taken to be the Cq representing the biological sample. The standard deviation (SD) of the triplicates was also calculated and samples that exhibited SD>0.20 were considered inconsistent. In such cases, one outlier Cq was removed to have at least duplicate Cq values for each biological sample and an SD<0.20.

For the DIV1 to DIV7 longitudinal dataset, reference gene validation was performed according to our qPCR data analysis workflow (Sundaram et al., 2019). Briefly, 10 conventional reference genes were chosen and screened using Coefficient of variation (CV) analysis and NormFinder (Supplementary Table S1) in R (https://moma.dk) (Andersen et al., 2004). The algorithm predicted *Tbp* and *Ppia* to be the most stable references. The normalization factor was then determined as the mean Cq value of *Tbp* and *Ppia* for each sample (Supplementary Table S1). For the comparison of DIV7 vs Schwann cell monocultures, reference gene validation was performed using CV analysis. *Mrpl10 and Sdha* exhibited the least collective variation (CV = 22%) and they were subsequently used for calculating the normalization factor. Relative expression of target genes was quantified using the 2^−ΔΔCt^ method (Livak and Schmittgen, 2001; Schmittgen and Livak, 2008) and data was visualized using Prism v8.0.

To assess statistical difference in relative RNA quantities between groups, One-way ANOVA was performed in Graph Pad Prism v8.0. If statistical significance was observed between the means of the groups, Tukey’s post hoc was performed to compare all the groups with each other. The alpha value threshold was set at 5% and the P-values are represented as follows: P<0.05 - *, P<0.01 - **, P<0.001 - ***.

## Results

### Progression of dissociated DRG/SC cocultures

We first documented the cellular composition of DRGs at E13.5 as well as the progression of the culture at Days *in Vitro* (DIV) 1, DIV3, DIV5 and DIV7 **(Figure 1)**. DRGs at E13.5 are comprised of sensory neurons with axonal projections towards the dorsal roots and the peripheral nerve. They are also comprised of neural crest derivatives (Sox10^+^ cells). Once dissected and dissociated, we observed a very sparse population of dissociated cells that comprised of neurons and neural crest derivatives (Sox10^+^ cells) at DIV1. At DIV3, however, we could see a neurite network gradually being formed with cells located on top of neuronal extensions. These cells are presumably migratory Schwann cell precursors. The neurites then grew out and established a well-connected network by DIV5. Schwann cells now densely populated neurites. Not much difference was observed between DIV5 and DIV7 except that more connections were established in the neurite-Schwann cell network **(Figure 1)**. From these observations, we could only deduce that Schwann cell precursors appear between DIV1 and DIV3 and they continue to populate the culture during the first 7 days.

### Expression profiles of Schwann cell developmental genes

We then determined the expression profiles of Schwann cell developmental genes using RTqPCR (**Figure 2A**). The genes that we assayed include *Dhh, Mpz, Mbp, Plp and Cnp* which are expressed in the Schwann cell lineage from the SCP stage *in vivo* (Jessen and Mirsky, 2019). We also assayed *Cad19* and *Tfap2α* which are expressed in SCP but are downregulated in iSC *in vivo* (Stewart et al., 2001; Takahashi and Osumi, 2005). Finally, we also assayed Krox20 mRNA levels which are upregulated as SCP transition into iSC in embryonic nerves (Topilko et al., 1994; Ghislain and Charnay, 2006)

**Figure 2:**
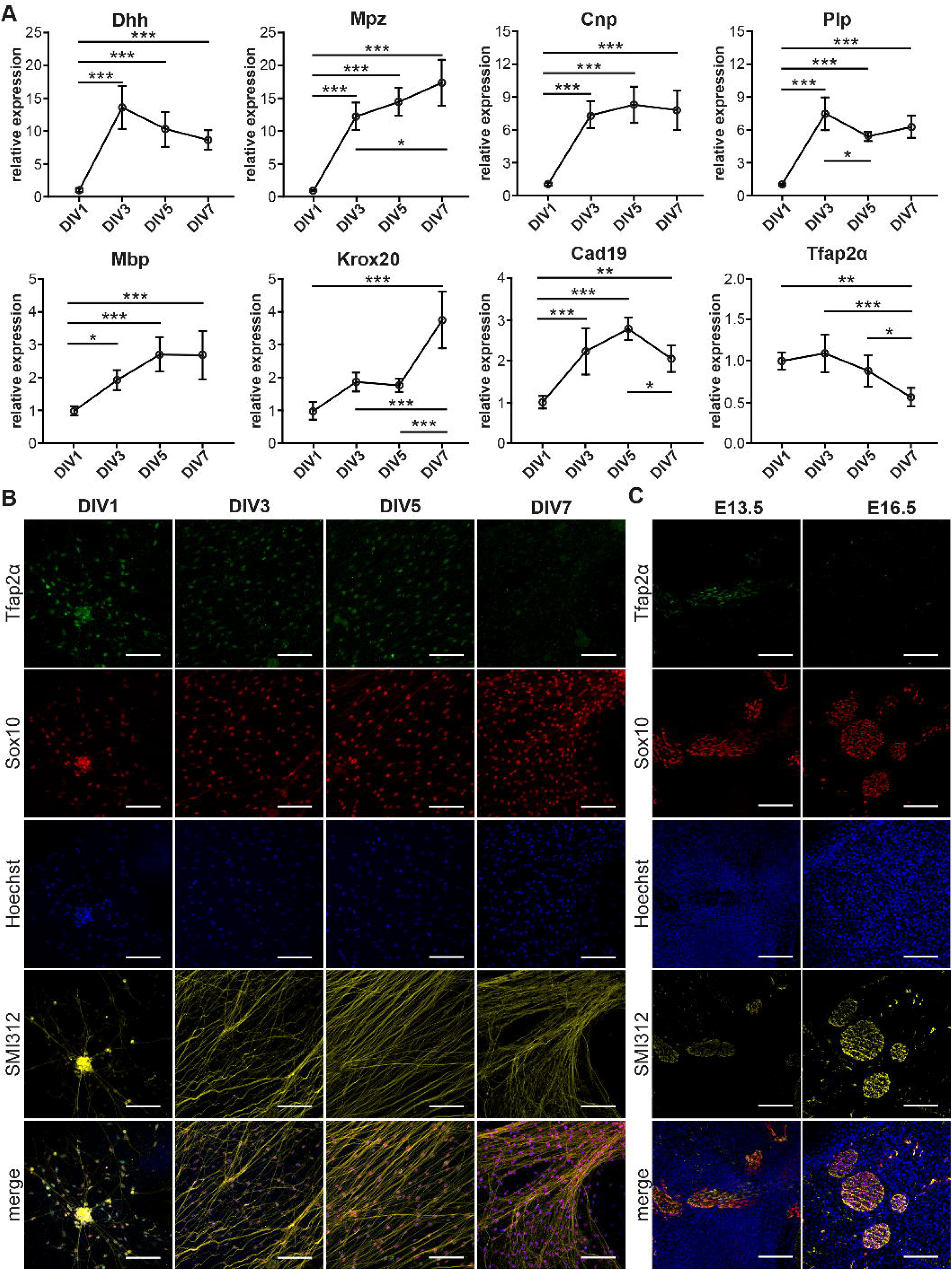
Transition of Schwann cell Precursors to immature Schwann cells in DRG/SC Cocultures. (A) mRNA expression profiles of Schwann cell lineage markers assessed through RTqPCR. Relative expression was calculated using DIV1 as the experimental calibrator. To assess statistical difference in relative RNA quantities between groups, One-way ANOVA was performed. If statistical significance was observed between the means of the groups, Tukey’s post hoc was performed to compare all the groups with each other. The alpha value threshold was set at 5% and the P-values are represented as follows: P<0.05 - *, P<0.01 - **, P<0.001 - ***. (B) ICC performed on dissociated cultures between DIV1 and DIV7. SCP were identified using Tfap2α staining, Sox10 was used as a SC lineage marker, Neurons were identified using SMI312 staining, and Hoechst dye was used to stain all nuclei. Scale bar = 100μm (C) IHC performed on hind limb cross sections of E13.5 and E16.5 embryos. SCP were identified using Tfap2α staining, Sox10 was used as a SC lineage marker, Neurons were identified using SMI312 staining, and Hoechst dye was used to stain all nuclei. Scale bar = 100μm

*Dhh* showed a stark 14-fold increase between DIV1 and DIV3. Although the profile seemed to project a downward trajectory after DIV3, the levels of *Dhh* did not vary significantly between DIV3 and DIV7. *Mpz* displayed an initial spike between DIV 1 and DIV3 by about 12 folds, which was similar to *Dhh*. However, the quantity of *Mpz* gradually increased and reached about 17 folds at DIV7. *Cnp* expression increased by about 7-folds between DIV1 and DIV3 and maintained a stable profile till DIV7. *Plp* showed an initial peak at DIV3 by about 7-folds, which was comparable to that of *Cnp*. However, *Plp* expression momentarily dropped at DIV5 and reached a plateau by DIV7. *Mbp* displayed a modest but statistically significant increase by 2-folds between DIV1 and DIV3. However, the expression did not increase significantly beyond DIV3. *Krox20* expression did not vary significantly between DIV1 to DIV5. However, at DIV7 we observed a sudden spike by about 4-folds. *Cad19* expression increased almost linearly between DIV1 and DIV5. However, between DIV5 and DIV7, we observe a drop in its expression level. *Tfap2α* maintains a flat profile from DIV1 and DIV3. It begins to decline after DIV3 and drops significantly to about 0.5 folds at DIV7.

### Expression of Tfap2α protein in Dissociated DRG/SC cocultures

The downregulation of Tfap2α protein levels in SCP is essential for their transition into iSC *in vivo* (Stewart et al., 2001). Consequently, we assayed the expression of Tfap2α in dissociated cultures using ICC **(Figure 2B)**. We observed the expression of the protein at DIV1, DIV3 and DIV5. However, at DIV7 we observed a huge reduction in Tfap2α immunoreactivity to background levels. This observation is also corroborated with the reduction in Tfap2α mRNA levels between DIV5 and DIV7 **(Figure 2A)**. The reduction in Tfap2α immunofluorescence is also observed *in vivo* wherein SCP at E13.5 in sciatic nerve transverse sections express the protein whereas iSC at E16.5 do not **(Figure 2C)**. These results collectively suggest that SCP in DRG/SC cocultures transition into iSC between DIV5 and DIV7.

### iSC/mSC transition in dissociated DRG/SC cocultures

We next sought to determine if iSC at DIV7 transition into pro-myelinating SC (pro-mSC) and myelinating SC (mSC) in cocultures upon the induction of myelination. In embryonic nerves, iSC differentiate into pro-mSC and mSC perinatally (E16.5 + around 4 days) (Salzer, 2015; Fledrich et al., 2019). These stages of Schwann cell development are characterized by the expression of Oct6 (pro-mSC) and Krox20 (mSC) transcription factors (Topilko et al., 1994; Jaegle et al., 1996, 2003). However, myelination *in vitro* requires the addition of Ascorbic Acid (AA) to cocultures to promote the formation of SC basal lamina and SC intrinsic epigenetic modifications which are prerequisites to promote further Schwann cell differentiation (Eldridge et al., 1987, 1989; Bacallao and Monje, 2015; Huff et al., 2020).

We treated cocultures at DIV7 with Ascorbic Acid and assayed Oct6 **(Figure 3A)** and Krox20 **(Figure 3B)** immunoreactivity after 4 days of treatment. We observed the presence of Oct6^+^ and Krox20^+^ Schwann cells located on top of the axons once the differentiation process is stimulated by AA addition. These results show that SC at DIV7 differentiate into pro-mSC and mSC after AA supplementation which is comparable to the perinatal iSC/mSC transition *in vivo*.

**Figure 3:**
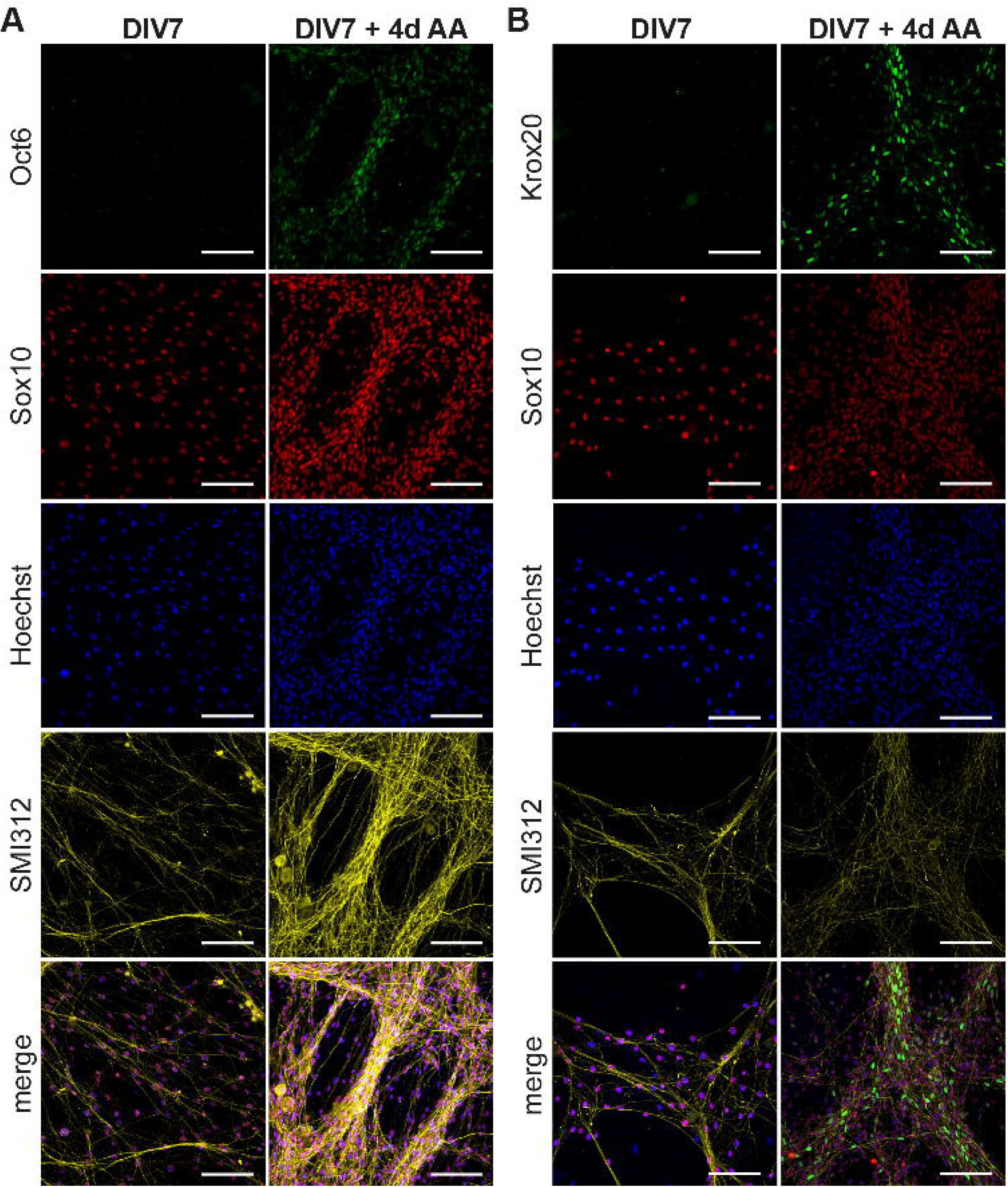
Transition of immature SC to myelinating SC in DRG/SC cocultures. Cocultures at DIV7 were treated with Ascorbic Acid (AA) for a period of 4 days. ICC was performed to assess the presence of Oct6+ (A) and Krox20+ (B) Schwann cells. Sox10 was used as a SC lineage marker, SMI312 was used to stain neurons and Hoechst dye was used to stain all nuclei. Scale bar = 100μm

### Myelinating potential of Coculture derived SC monocultures

SC monocultures are established from enzymatic digestion of neonatal mouse sciatic nerves in a plethora of recent studies because the preparation technique is less cumbersome. Neonatal nerves contain underdeveloped connective tissues and unmyelinated fibers and can be easily digested to render copious amounts of primary SC (Monje, 2020). Myelination can be induced in these cells by the addition of cAMP in substantially large concentrations (Arthur-Farraj et al., 2011; Bacallao and Monje, 2015). We therefore wanted to investigate if primary SC monocultures established from DIV7 cocultures are comparable to cultures established from neonates. We isolated SC from DIV7 cocultures using immunopanning and expanded them in culture for 48h (Proliferation) following which these cells were treated with cAMP for another 48h (Differentiation) according to prescribed protocols employing neonatal SC cultures (Arthur-Farraj et al., 2011). We assessed for the presence of pro-mSC and mSC using Oct6 and Krox20 mRNA and protein expression using DIV7 cells as experimental controls **(Figure 4)**.

**Figure 4:**
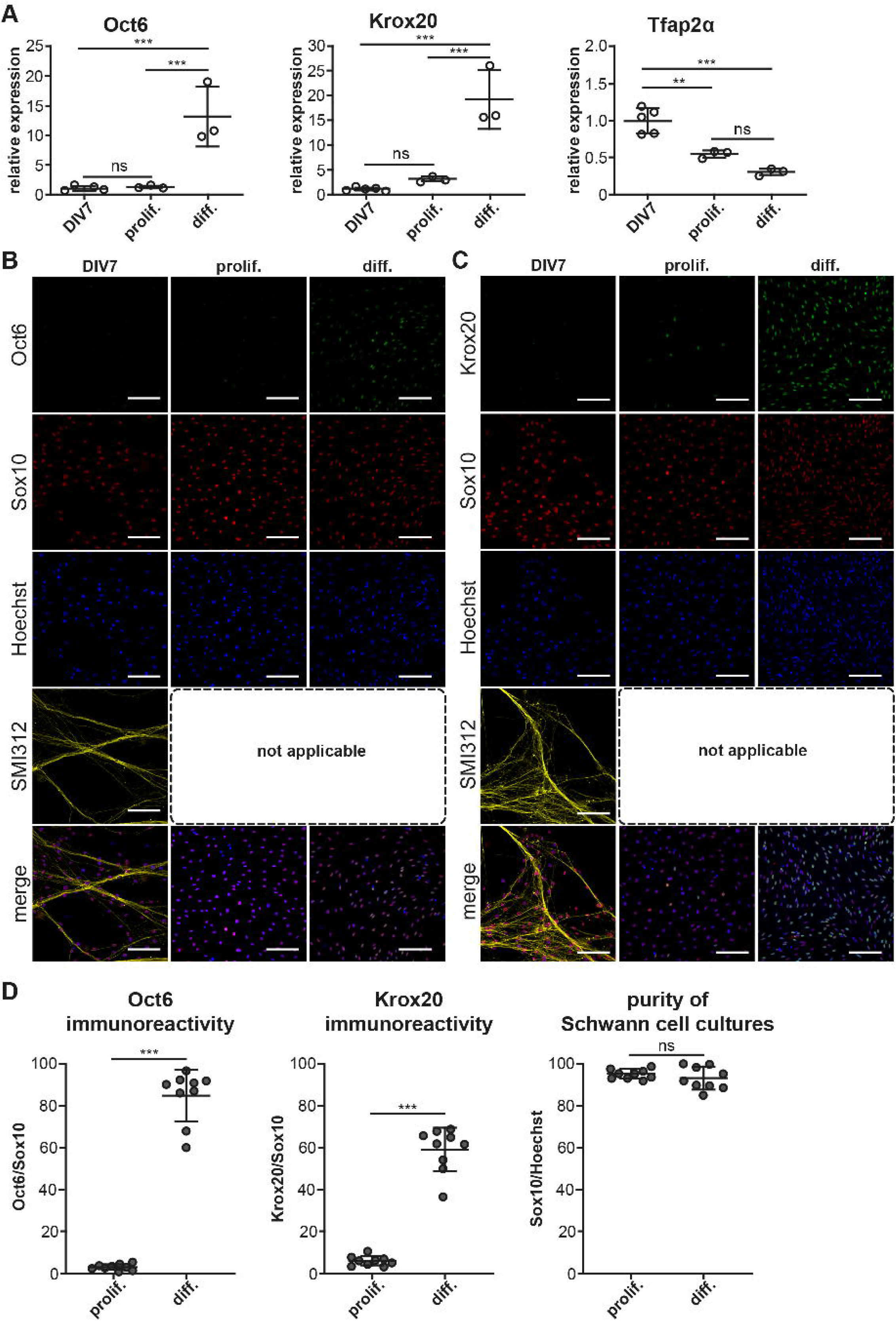
Myelinating potential of Coculture derived SC monocultures. SC monocultures were established from DIV7 DRG/SC cocultures using immunopanning and were differentiated using cAMP. (A) Relative mRNA of Oct6, Krox20 and Tfap2α assessed across DIV7, SC monocultures in Proliferation (Prolif.) and in Differentiation (Diff.) DIV7 was used as the experimental control. To assess statistical difference in relative RNA quantities between groups, One-way ANOVA was performed. If statistical significance was observed between the means of the groups, Tukey’s post hoc was performed to compare all the groups with each other. The alpha value threshold was set at 5% and the P-values are represented as follows: P<0.05 - *, P<0.01 - **, P<0.001 - ***. (B) ICC performed on dissociated cultures at DIV7 as well as in SC monocultures in proliferation and in differentiation. Pro-mSC were identified using Oct6 staining, Sox10 was used as a SC lineage marker, Neurons were identified using SMI312 staining, and Hoechst dye was used to stain all nuclei. SMI312 is not applicable in SC monocultures due to the absence of axons. Scale bar = 100μm (C) ICC performed on dissociated cultures at DIV7 as well as in SC monocultures in proliferation and in differentiation. mSC were identified using Krox20 staining, Sox10 was used as a SC lineage marker, Neurons were identified using SMI312 staining, and Hoechst dye was used to stain all nuclei. SMI312 is not applicable in SC monocultures due to the absence of axons. Scale bar = 100μm (D) Quantification of Sox10, Oct6 and Krox20 immunoreactivity. The purity of SC cultures was assessed by calculating the total number of Sox10+ cells in culture. Oct6 and Krox20 immunoreactivity was quantified as the ratio of the total number of Oct6+ and Krox20+ cells to Sox10+ cells and it is expressed as a percentage. To assess statistical differences, a non-parametric Mann Whitney test was performed. The alpha value threshold was set at 5% and the P-values are represented as follows: P<0.05 - *, P<0.01 - **, P<0.001 - ***.

mRNA expression levels of Oct6 and Krox20 do not vary much between DIV7 dissociated DRG/SC coculture and SC monocultures in proliferation medium but we noticed a stark increase in Oct6 mRNA levels by about 12 folds and Krox20 mRNA by about 18 folds when SC monocultures were treated with cAMP to induce differentiation **(Figure 4A)**. This increased mRNA levels also corresponded with an increase in Oct6 and Krox20 immunoreactivity **(Figure 4B, 4C, quantified in 4D)** where we observed about 80% of Sox10^+^ SC also stained positive for Oct6 and about 60% of SC stained positive for Krox20 on an average. Very similar results are also observed in SC monocultures derived from neonatal sciatic nerves (Arthur-Farraj et al., 2011)

Furthermore, we also inspected the mRNA and protein levels of the SCP marker Tfap2α. Interestingly, Tfap2α mRNA levels are further downregulated by about 0.5 folds between DIV7 dissociated DRG/SC coculture and Proliferative SC monocultures **(Figure 4A)**. Besides, we could not detect any Tfap2α^+^ cells in all the three conditions (Supplemental Figure S2). These results suggest that SC monocultures established from DIV7 cocultures are comparable to primary cultures established from neonatal sciatic nerves with regard to the change in SC phenotype upon cAMP addition. Furthermore, we also observed that these monocultures do not express the SCP marker Tfap2α at the protein level. Taken together, these results provide further evidence that DIV7 SC are phenotypically different from SCP that arise in DRG/SC cocultures between DIV1 and DIV5 and they are rather similar to neonatal SC in culture.

## Discussion

The objective of this study was to determine the SCP/iSC/mSC transition in DRG/SC cocultures using the expression profiles of Schwann cell developmental genes (*Dhh, Mpz, Cnp, Plp, Mbp, Krox20, Cad19, Tfap2α*). We first determined the expression profiles of these genes between DIV1 and DIV7. In addition to the mRNA data, we used Tfap2α ICC as a confirmatory experiment to distinguish between SCP and iSC in the coculture model. Furthermore, we induced myelination by the addition of Ascorbic Acid at DIV7 to verify that the iSC transition into pro-mSC ad mSC *in vitro*. Additionally, we also compared the myelinating potential of SC monocultures established from DIV7 cocultures to that of monocultures established from neonatal nerves. A detailed schematic of our experimental approach is detailed in **Figure 5A**.

**Figure 5:**
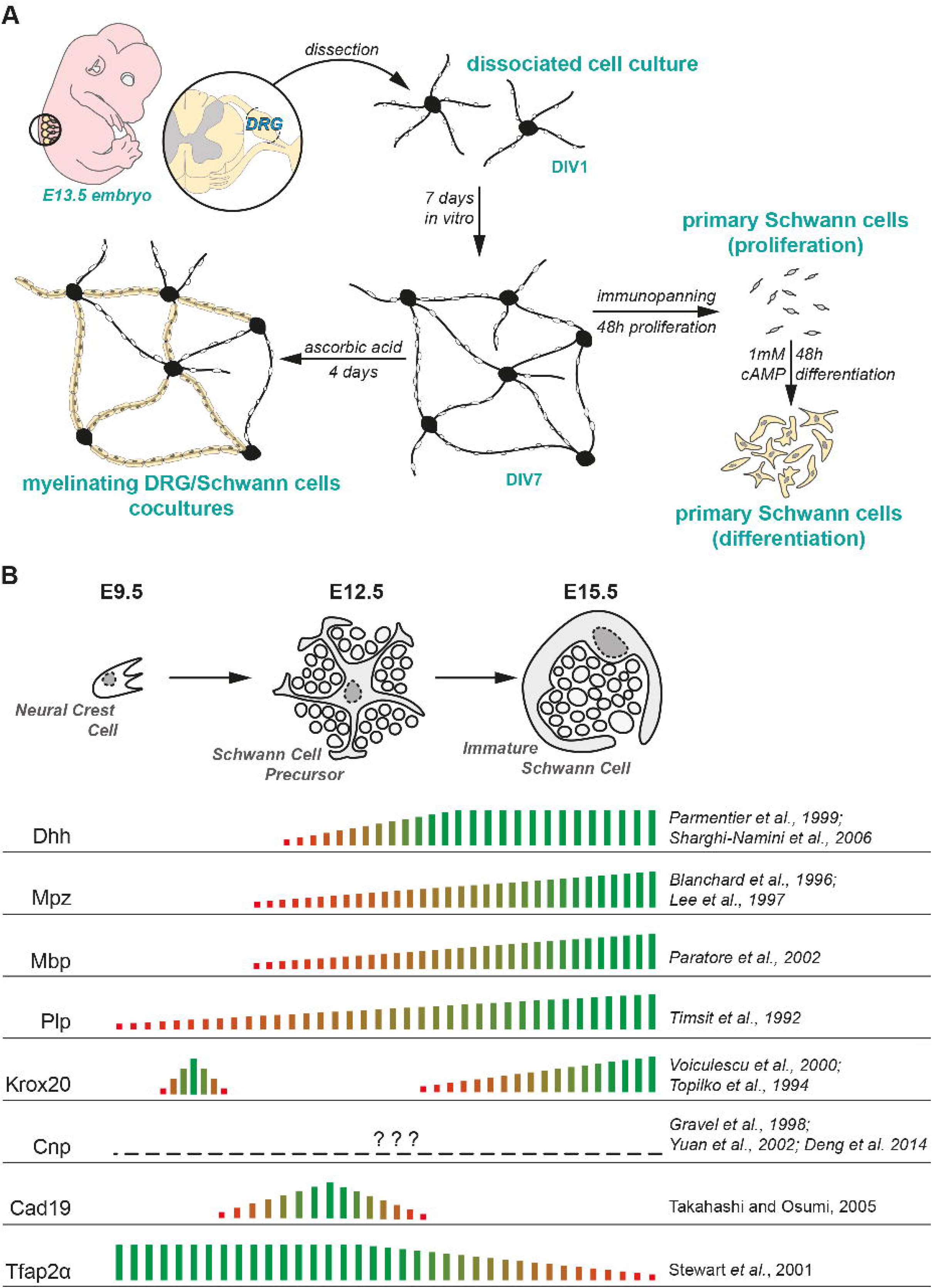
Experimental paradigm and literature review. (A) Experimental design used in the study. (B) Current consensus on the mRNA expression of SC lineage markers in embryonic peripheral nerves presented along with the corresponding literature. The empirical quantification presented herein has been extrapolated from microarray studies performed on murine embryonic nerves (Buchstaller et al., 2004; D’Antonio et al., 2006).

### The SCP/iSC transition

The expression profiles of SC lineage genes have been previously documented in gene profiling studies on murine embryonic nerves (Buchstaller et al., 2004; D’Antonio et al., 2006). Seminal reviews on Schwann cell developmental markers and other gene-specific expression profiling studies have been published in the last couple of decades and they give us a holistic understanding of the SC developmental transitions in embryonic and postnatal peripheral nerves (Jessen et al., 1994; Jessen and Mirsky, 2005; Woodhoo and Sommer, 2008; Monk et al., 2015). A summary of the current consensus on the expression profiles of these genes *in vivo* along with the relevant literature is presented in **Figure 5B**.

When comparing our mRNA expression data to these profiles, we find that our data seem to largely comply and concur with the order of molecular events observed in embryonic spinal nerves. For instance, the expression of *Dhh, Mpz, Mbp & Plp* is upregulated in SCP at around E12.5 in mice (Jessen and Mirsky, 2005; Woodhoo and Sommer, 2008). In DRG cultures, *Dhh, Mpz, Mbp and Plp* expression is increased at DIV3 (**Figure 2A**). These observations suggest that SC in coculture either increase their endogenous expression of these SCP markers or that they proliferate extensively between DIV1 and DIV3. We find that the latter is more plausible as SC in cocultures continue to proliferate from DIV1 all through DIV7 (*Supplementary Figures, Figure S1 for Ki67 staining in cocultures*). Moreover, it is worth noting that the cells have already attained the SCP state at the time of dissection (E13.5) (Jacob, 2015).

Data on *Cnp* expression in SCP of embryonic nerves is inconclusive. We analyzed three separate studies that sought to trace *Cnp* expression in the PNS (Gravel et al., 1998; Yuan et al., 2002; Deng et al., 2014). One study demonstrated that *Cnp* is expressed in the satellite cells of DRGs at E14.5 and at the ventral roots at E17.5 but did not comment on its expression in SCPs (Deng et al., 2014). The other two studies dealt with post-natal time points. Nonetheless, our observations suggest that *Cnp* mRNA is upregulated in SCP from DIV3 in culture. This observation, however, needs to be verified *in vivo*.

*Krox20* is expressed at two different time points during the development of the PNS (Topilko et al., 1994; Voiculescu et al., 2000; Coulpier et al., 2010). At around E10.5, it is first expressed in the boundary cap cells that are located at the dorsal and ventral roots. This is followed by its increased expression as SCP transition into iSC at around E15.5 in peripheral nerves. Indeed, it is one of the genes used in the study that can categorically distinguish between SCP and iSC at the mRNA expression level. In our study, Krox20 expression does not change significantly between DIV1 and DIV5. We then see a sudden spike in its expression between DIV5 and DIV7 suggesting that SCP transition to iSC in this time-period (**Figure 2A**).

To test this hypothesis, we further looked at the expression profile of *Tfap2α*, which is expressed in NCC and SCP but not in iSC (Stewart et al., 2001) (**Figure 2A**). The downregulation of this transcription factor is indeed required for SCP to transition into iSC in developing nerves (Stewart et al., 2001; Jacob, 2015). Consistent with this observation, *Tfap2α* levels drop by 50% between DIV5 and DIV7 thus providing further evidence of the SCP/iSC transition between DIV5 and DIV7. Furthermore, Tfap2α immunorecativity in dissociated DRG/SC cocultures starkly reduces at DIV7 **(Figure 2B)** congruent with its downregulated expression in iSC *in vivo* **(Figure 2C)**. Finally, we also assayed the expression levels of *Cad19*, which is the only known gene that is uniquely expressed in SCP but neither in Neural Crest Cells nor in iSC (Jessen and Mirsky, 2005, 2019; Takahashi and Osumi, 2005). *Cad19* expression *in vitro* was the highest around DIV3 and DIV5 but it reduced again at DIV7 reaching to levels comparable to DIV3. It is to be noted that all these changes in mRNA expression occur despite continuous proliferation of SC in this time period (Supplementary Data, Figure S1) suggesting that cells indeed downregulate *Cad19* and *Tfap2α* between DIV5 and DIV7.

Taken together, these results give a clear picture of Schwann cell developmental transitions in embryonic cocultures during which SCP proliferate and also presumably migrate on developing neurites between DIV1 and DIV5. At DIV5, SCP begin their transition into iSC and at DIV7 most of the cells present on neurites are iSC.

It is interesting to note that SCP monocultures established directly from E12.5 dissociated mouse peripheral nerves transition into iSC in cultures after 4 DIV (E12.5 + 4DIV) which corresponds exactly to their timing *in vivo* (Dong et al., 1999). However, it takes up to 7 DIV to achieve this transition in DRG/SC cocultures. One plausible reason for this difference is the fact that SCP monocultures are expanded in the presence of a fixed concentration of Neuregulin to mimic the trophic support from axons that in turn drives their differentiation to iSC (Dong et al., 1995; Leimeroth et al., 2002). In cocultures however, the media is not supplemented with Neuregulin and neurites have to first emanate from the dissociated soma and achieve significant growth during the first 72h. Therefore, it is possible that optimal levels of axonal Neuregulin are not present in the coculture system until DIV5 thereby causing a delay in the SCP/iSC transition. This hypothesis can also be extended to other axo-glial signaling pathways such as Notch signaling which is also crucial for the SCP/iSC transition (Woodhoo et al., 2009). In effect, we think that the delay in the SCP/iSC transition in cocultures is largely driven by the fact that sufficient axonal extensions (and therefore axonal differentiation and axonal cues) are only achieved between DIV5 and DIV7. However, this hypothesis warrants further experimentation.

### The iSC/mSC transition

iSC *in vivo* give rise to promyelinating and myelinating SC perinatally which is about 4 days after iSC emante from SCP (Salzer, 2015; Fledrich et al., 2019). Therefore, we hypothesized that if the cells at DIV7 are indeed iSC, then they should transition into pro-mSC and mSC upon the addition of Ascorbic Acid. We were able to observe Oct6^+^ and Krox20^+^ cells after 4 days of Ascorbic Acid (AA) supplementation **(Figure 3)**. However, Oct6 expression *in vivo* is only transient, and it promotes the differentiation of pro-mSC to mSC (Jaegle et al., 1996, 2003). In other words, perinatal SC *in vivo* sequentially express Oct6 and Krox20 but fully mature myelinating SC do not express Oct6 postnatally.

Our data show that both these cellular phenotypes are present after 4 days of AA supplementation suggesting that iSC/mSC transition is still ongoing at this timepoint. It is also equally possible that the cells that express Krox20 at DIV7 + 4 days AA expressed Oct6 before and they have already transitioned to mSC. Therefore, further experiments are required to assess the advent and the successive extinction of Oct6 expression in this coculture system. Nevertheless, our results clearly demonstrate that iSC at DIV7 transition into mSC as early as after 4 days of AA supplementation which is comparable to their transition *in vivo*.

As for SC monocultures, the phenotype of SC in culture; irrespective of the source, is highly peculiar as these cells express several makers of the Schwann cell lineage including those of SCP, iSC and the repair phenotype that arises post nerve injury (Monje, 2020). In our study, we wanted to determine the myelination potency of coculture derived SC monocultures and compare that to SC monocultures established from neonatal nerves which largely comprise of iSC and pro-mSC (Monje, 2020) **(Figure 4)**. Our observations demonstrate firstly that SC monocultures derived from DIV7 cocultures are similar to cultures established from neonatal peripheral nerves in their potential to differentiate into mSC. Secondly, and more importantly, they show that DIV7 SC in monoculture are distinct from SCP that arise between DIV1 and DIV5 in cocultures as they lack Tfap2α immunoreactivity both in proliferative and in differentiative conditions (Supplementary figure S2). However, we have not investigated any fine phenotypic differences between neonatal SC monocultures and coculture-derived SC monocultures as it falls beyond the ambit of the present study. Nevertheless, it is an important criterion that remains to be ascertained. That being said, it would also be equally interesting to isolate SCP from cocultures before DIV5 and compare them to SCP derived directly from embryonic nerves at E12.5 to ascertain phenotypic differences, if any.

In summation and as highlighted above, our study demonstrates for the first time that the SCP/iSC transition indeed occurs in embryonic DRG/SC cocultures between DIV5 and DIV7. The further differentiation of iSC to mSC can also be observed in this model as early as 4 days post AA supplementation. These observations serve as a powerful frame of reference to design, execute and comprehend coculture experiments using different Schwann cell mutants that aim to ascertain the role of specific genes or experimental conditions in Schwann cell development and myelination.

## Acknowledgements

The authors thank the *Animal House Core Facility* and the *Cyto2BM Molecular Biology Platform* of BioMedTech Facilities (INSERM US36/CNRS UMS2009) for the animals and research services pertaining to the generation of qPCR data. The authors also thank the SCM microscopy platform for the services pertaining to confocal imaging. The authors would also like to thank Prof. Dies MEIJER, University of Edinburgh, for the Krox20 and Oct6 antibodies. We also thank Prof. Rhona MIRSKY and Prof. Kristjan R JESSEN, University College London, for the critical reading of the preprint and for their inputs.

