## Supplementary Figures for "Retracing Schwann cell developmental transitions in embryonic dissociated DRG/Schwann cell cocultures in mice"

**Figure S1: Ki67 ICC DIV1 – DIV7**

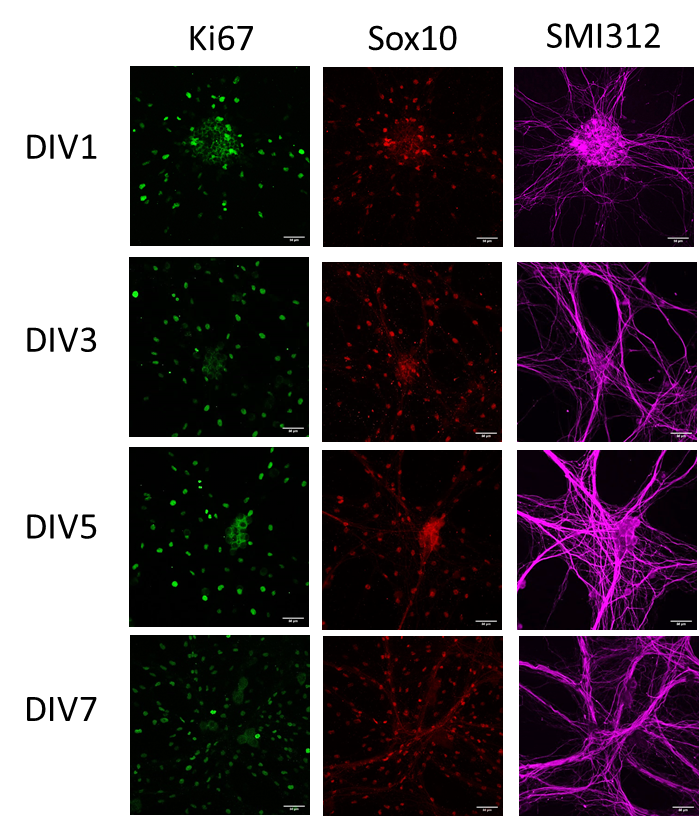

Sox10+ SC are also Ki67+ across all DIVs suggesting that SC continue to proliferate even at DIV7.

**Figure S2: Tfap2α ICC DIV7 vs Prolif. vs Diff**

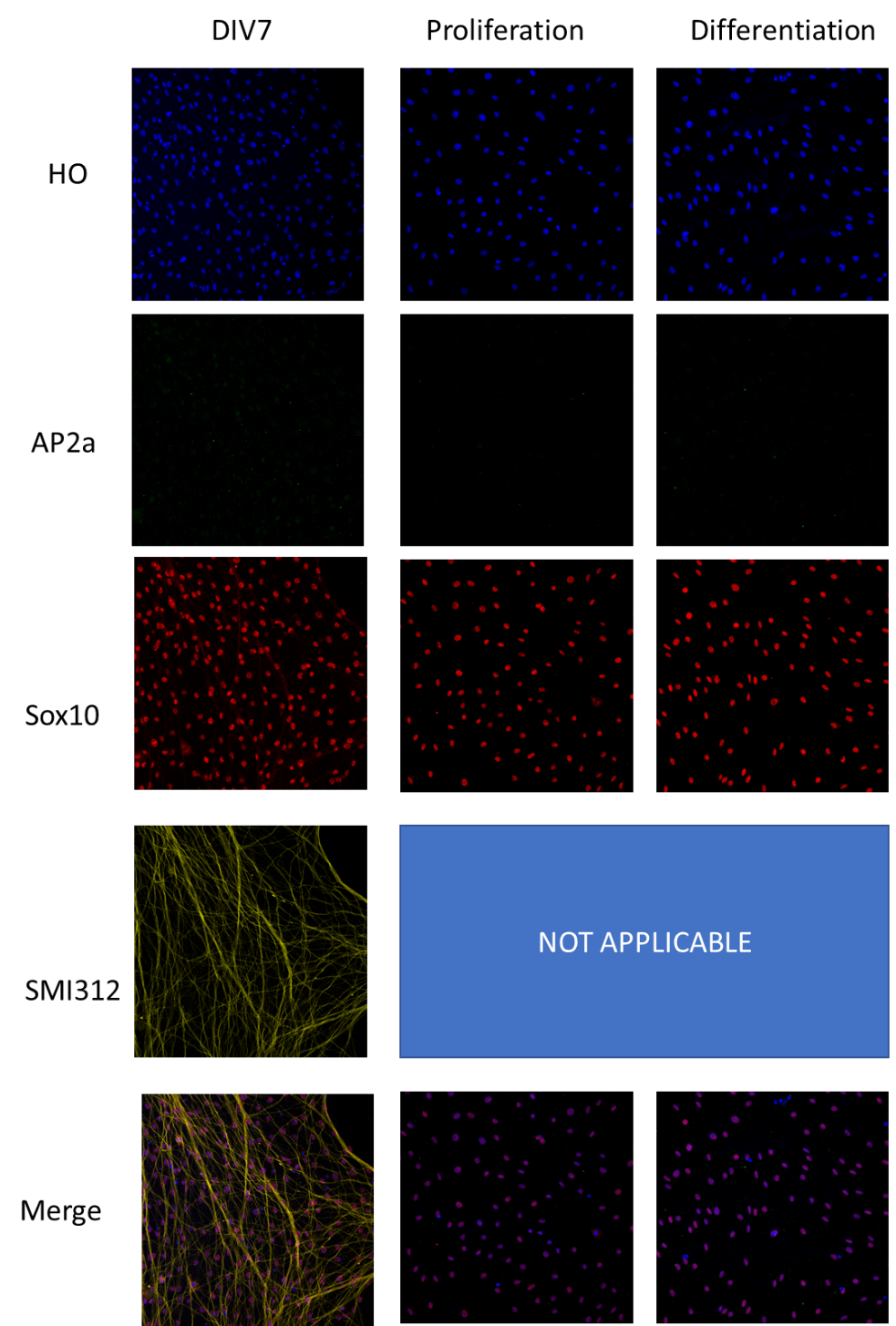

Tfap2α immunoreactivity not detected in any of the three conditions suggesting that cells at DIV7 and in monocultures are not SCP.
