## Supplementary Methods for "Retracing Schwann cell developmental transitions in embryonic dissociated DRG/Schwann cell cocultures in mice"

**Media compositions:**

**DRG Plating Media**

| **Component** | **Final Concentration** | **Reference** |
| --- | --- | --- |
| DMEM high glucose | Base medium | 11960044 Gibco |
| Glutamax (100X) | 1X | 35050061 Gibco |
| Heat inactivated Horse Serum | 10% | 26050088 Gibco |
| 2.5S NGF | 50ng/ml | 1156NG R&D |
| Antibiotic/ Antimycotic (100X) | 1X | 15240062 Gibco |

**Neurobasal Media**

| **Component** | **Final Concentration** | **Reference** |
| --- | --- | --- |
| Neurobasal Medium | Base medium | 21103049 Gibco |
| Glutamax (100X) | 1X | 35050061 Gibco |
| 2.5S NGF | 50ng/ml | 1156 NG R&D |
| B27 supplement (50X) | 1X | 17504044 Gibco |
| Antibiotic/ Antimycotic (100X) | 1X | 15240062 Gibco |

Ascorbic Acid (A4403 Sigma) was added to DRG plating media at a final concentration of 50µg/mL to induce myelination in cocultures.

Coverslips were coated with PLL (P7890, Sigma) at a final concentration of 0.05mg/mL in 0.15M Sodium Borate Buffer for 30 mins. Coverslips were washed and further coated with Collagen (3440-100-01, R&D system, 1:7 final dilution) in the presence of 35% Ammonia for 15 min. Collagen was removed leaving behind a thin layer upon which the cells were plated.

Immunopanning was performed as described in Lutz A.B., CSH Protocols, 2014. A detailed protocol adapted to DRG/SC cocultures can be made available upon request.

**Schwann cell Proliferation Media:**

| **Component** | **Final Concentration** | **Reference** |
| --- | --- | --- |
| DMEM: F12 1:1 | Base medium | 21331046 Gibco |
| Glutamax (100X) | 1X | 35050061 Gibco |
| N2 supplement (100X) | 1X | 17502-048 Gibco |
| Neuregulin | 10ng/ml | 396 HB 050 R&D |
| Forskolin | 2.5µM | F3917 Sigma |
| Antibiotic/ Antimycotic (100X) | 1X | 15240062 Gibco |
| B27 supplement (50X) | 1X | 17504044 Gibco |
| T3 salt | 20 ng/µl | T6397 Sigma |

**Schwann cell Differentiation Media:**

| **Component** | **Final Concentration** | **Reference** |
| --- | --- | --- |
| DMEM: F12 1:1 | Base medium | 21331046 Gibco |
| Glutamax (100X) | 1X | 35050061 Gibco |
| N2 supplement (100X) | 1X | 17502-048 Gibco |
| Neuregulin | 10ng/ml | 396 HB 050 R&D |
| dbcAMP | 1mM | D0627 Sigma |
| Antibiotic/ Antimycotic (100X) | 1X | 15240062 Gibco |
| B27 supplement (50X) | 1X | 17504044 Gibco |
| T3 salt | 20 ng/µl | T6397 Sigma |

**Antibodies and Concentrations:**

| **I Ab** | | | **II Ab** | | |
| --- | --- | --- | --- | --- | --- |
| Protein | Ref | Conc | Type | Ref | Conc |
| **Sox10** | AF2864 (R&D) | 1/200 | **Donkey anti Goat Cy3** | 705-165-147 (Jackson) | 1/500 |
| **Tfap2α** | ab108311 (Abcam) | 1/100 | **Donkey anti Rabbit A488** | 711-545-152 (Jackson) | 1/200 |
| **Krox20** | Gift from Prof. Meijer | 1/100 | **Donkey anti Rabbit A488** | 711-545-152 (Jackson) | 1/200 |
| **Ki67** | ab15580 (Abcam) | 1/1000 | **Donkey anti Rabbit A488** | 711-545-152 (Jackson) | 1/1000 |
| **Oct6** | Gift from Prof. Meijer | 1/100 | **Donkey anti Rabbit A488** | 711-545-152 (Jackson) | 1/200 |
| **SMI312** | 837904 (Biolegend) | 1/250 | **Donkey anti Mouse DyLight 633** | NBP1-75613 (Novusbio) | 1/500 |

Normal Donkey Serum: Ref 017-000-121 (Jackson)

**Primer sequences:**

| Refer to Sundaram et al, 2019, Plos One, for the primer sequences of reference genes | | |
| --- | --- | --- |
| The sequences for the target genes are given below | |  |
| **Primer Name** | **Sequence** | **Efficiency** |
| m_Dhh_F | ATCCACGTATCGGTCAAAGC | 100% |
| m_Dhh_R | AGTCACCACGATGTAGTTCCC |  |
| M_Mpz F | ATCTCTTTTACCTGGCGCTACC | 99.50% |
| M_Mpz R | ACTGGATGCGCTCTTTGAAG |  |
| M_Mbp F | ACTCACACACGAGAACTACCC | 99% |
| M_Mbp R | GGTGTTCGAGGTGTCACAATG |  |
| m_Krox20 F | TGCACCTAGAAACCAGACCTTC | 95% |
| m_Krox20 R | TGCCCGCACTCACAATATTG |  |
| m_CNPase F | GACAGCGTGGCGACTAGACT | 96% |
| m_CNPase R | CACCTGGAGGTCTCTTTCCA |  |
| m_PLP F | AGCAAAGTCAGCCGCAAAAC | 98% |
| m_PLP R | CCAGGGAAGCAAAGGGGG |  |
| m_Tfap2a F | ACTCCTTACCTCACGCCATC | 97.50% |
| m_Tfap2a R | AGCATTGCTGTTGGACTTGG |  |
| m_Cad19 F | TGACATAGGGGAGAATGCAGAG | 99.50% |
| m_Cad19 R | AAGCTCTTCATCCACATGGC |  |
